# Bioelectric-calcineurin signaling module regulates allometric growth and size of the zebrafish fin

**DOI:** 10.1101/258442

**Authors:** Jacob M Daane, Jennifer Lanni, Ina Rothenberg, Guiscard Seebohm, Charles W Higdon, Stephen L Johnson, Matthew P Harris

## Abstract

The establishment of relative size of organs and structures is paramount for attaining final form and function of an organism. Importantly, variation in the proportions of structures frequently underlies adaptive change in morphology in evolution and maybe a common mechanism underlying selection. However, the mechanism by which growth is integrated within tissues during development to achieve proper proportionality is poorly understood. We have shown that signaling by potassium channels mediates coordinated size regulation in zebrafish fins. Recently, calcineurin inhibitors were shown to elicit changes in zebrafish fin allometry as well. Here, we identify the potassium channel *kcnk5b* as a key player in integrating calcineurin’s growth effects, in part through regulation of the cytoplasmic C-terminus of the channel. We propose that the interaction between Kcnk5b and calcineurin acts as a signaling node to regulate allometric growth. Importantly, we find that this regulation is epistatic to inherent mechanisms instructing overall size as inhibition of calcineurin is able to bypass genetic instruction of size as seen in *sof* and wild-type fins, however, it is not sufficient to re-specify positional memory of size of the fin. These findings integrate classic signaling mediators such as calcineurin with ion channel function in the regulation of size and proportion during growth.

## Introduction

The establishment of relative proportion of structures and organs is essential for the normal physiology and function of an organism. The study of differential growth of structures in development and in the evolution of form has a rich history (Gayon, 2000). A key advance in our understanding differential growth stems from the work of D’Arcy Thompson and his efforts to define the underlying rules of coordinated transformations in form (Thompson, 1917). This foundational work was leveraged by Huxley and Teissier who formalized scaling relationships between structures within organisms as a power law that details the relative proportion of structures (Huxley and Teissier, 1936; Julian Huxley, 1932)). This law provides a scale-independent means of comparing growth in development and among species.

A common mechanism for the establishment of organ size and the relative proportions of structures is through differential sensitivity of organs to a systemic growth signal such as insulin-like growth factors (IGF) or growth hormone (GH) (Bryant and Simpson, 1984; Conlon and Raff, 1999). However, there is substantial evidence for an organ-intrinsic capacity to establish relative size that is robust to such broad systemic signals. Transplanted organs will frequently reach their target size even when cultured in ectopic locations (Dittmer et al., 1974; Felts, 1959; Twitty et al., 1931). Intrinsic regulation of size in structures is also observed in cases in which organs will accelerate growth back to a seemingly pre-ordained growth trajectory following growth-limiting insults, such as nutrient deprivation or illness (Finkielstain et al., 2013; Prader et al., 1963). This “catch-up growth” is also seen during epimorphic regeneration, wherein growth rate is dependent on the amount of tissue lost such that recovery of the original form occurs within the same time window; this suggests that relative growth rates are guided by *retained* positional cues within tissues (Lee et al., 2005; Morgan, 1906; Spallanzani, 1769; Tassava and Goss, 1966). How growth is integrated with positional information within the organ to achieve proper proportion remains unclear.

An effective strategy to understand growth and regulation of size is to analyze mutants or experimental conditions in which scaling properties have been altered. Through genetic screens in the zebrafish, several mutants have been identified that have adult fins that are reduced in size as well as mutants in which fins grow beyond their normal limit (Eeden et al., 1996; Fisher et al., 2003; Goldsmith et al., 2003; Green et al., 2009; Huang et al., 2009; Iovine and Johnson, 2000; Iovine et al., 2005; Perathoner et al., 2014). The *another longfin* mutant was identified as having enlarged fins and barbels (Haffter et al., 1996). We have previously shown that *alf* is caused by a specific gain-of-function point mutation in the potassium channel *kcnk5b*, a member of the TWIK/TASK two pore family (K_2P_) of potassium channels (Perathoner et al., 2014). Supporting a role for ion regulation in establishing proportion, the *shortfin* (*sof*) mutant has been mapped back to mutations in the gap junction *connexin43* (*cx43/gja1*) (Iovine et al., 2005; Sims et al., 2009). Though both gap junctions and potassium channels have important roles in physiology and cell biology, these initial findings in the zebrafish fin are surprising in that they uncover a specific role for local bioelectric signaling in the regulation and coordination of growth.

Insight into the mechanisms regulating coordinated growth also comes from pharmacological regulation of cell signaling during development. Inhibition of calcineurin in developing and regenerating fins by FK506 and cyclosporin A specifically increases the size of the resulting fin (Kujawski et al., 2014). Calcineurin is a protein phosphatase known to affect transcription through direct binding and dephosphorylation of the NFAT family of transcription factors (Hogan et al., 2003). Analysis of the patterning and gene expression profiles in overgrown fins treated with FK506 led Kujawski and colleagues to suggest that positional information of the fin was specifically altered by calcineurin inhibition. Under this hypothesis, activation of calcineurin triggers a proximal-like growth program that leads to enhanced proliferation to re-specify larger fin sizes (Kujawski et al., 2014).

Here, we extend analysis into mechanisms of size regulation by potassium channels and calcineurin using the zebrafish fin as a model. We show that FK506 can mirror the growth effects of activated Kcnk5b, and can override genetic determination of size. Importantly, we demonstrate that Kcnk5b function is a key component to the ability of FK506 to regulate growth. Through our genetic and experimental analyses, we demonstrate that the growth-mediating properties of FK506 do not re-specify positional memory in the fin, but rather act to maintain heightened rate of growth during regeneration of the fin. These data advance our understanding of how size is regulated during growth and provide a regulatory link between bioelectric and classical signaling pathways in the regulation of growth and proportion.

## Results

### Skeletal phenotypes of fins treated with FK506 compared to mutants affecting fin proportion

Fins are composed of multiple segmented rays of dermal bone. Elongation of the fin occurs through sequential addition of these bony segments to the distal fin tip. The zebrafish fin-ray segments are regularly patterned such that each segment is of roughly same length along the proximal-distal axis. While the segmentation pattern of the fin rays can be dissociated from total fin length (Schulte et al., 2011), mutants with altered fin size are often accompanied by a change in size of segments, consistent with alterations in the rate of growth (Goldsmith et al., 2003; Perathoner et al., 2014; Sims et al., 2009). The fin segments in *kcnk5b*/*alf* mutant are generally elongated, while those of the *shortfin* mutant are shorter (Iovine et al., 2005; Perathoner et al., 2014)(Figure S1).

As segmentation patterns are influenced by mutations that alter fin size, we asked whether FK506 treatment had an effect on segmentation patterning. We focused primarily on fin regeneration, which recapitulates growth properties that occur during fin development. During fin regeneration, both the missing fin tissue and segmentation patterns are precisely restored. Thus, regeneration assays can serve as a foundation for exploring the genetic contributions of positional identity and memory, growth rate, and patterning. Treatment of regenerating fins with the calcineurin inhibitors FK506 and cyclosporin A led to dose-dependent coordinated overgrowth of fin regenerates (Kujawski et al., 2014)(Figure S1E). FK506-treated fins also exhibited elongated fin ray segments, similar to those observed in *kcnk5b/alf* mutants (Figure S1E F,G). Surprisingly, FK506 treatment of *cx43/shortfin* regenerating fins also resulted in elongated fin ray segments (Figure S1H).

### FK506 treatment regulates growth independently of memory of size

As treatment of fins with FK506 appears to bypass the genetic specification of normal fin length in wild-type fish (Kujawski et al., 2014), we sought to define the growth characteristics of inhibiting calcineurin in the background of different mutants with altered fin size. Similar to wildtype, when cut to 50% of their initial size, the fins of short-finned (*gja1/sof*) and long-finned (*kcnk5b/alf)* fish regenerate back to their pre-amputation size over a similar time period despite their vastly different starting lengths (Figure 1). However, FK506-treatment of regenerating fins from both wild-type and *gja1*short-finned mutants leads to an increased rate of growth in each group, resulting in the formation of similarly sized, larger fins in both genotypes (Figure 1). Oddly, the shape and size of FK506-treated fins from *cx43/sof* mutants were indistinguishable from treated wild-type fins (Figure S1H). Of note, FK506-treatment of fish with gain-of-function of *kcnk5b* activity does not lead to an additional increase in the size of the regenerate or rate of its growth over untreated mutants (Figure 1). All fins treated with FK506 regrow at comparably increased rates, regardless of genotype or previous size. These data suggest that the effect of calcineurin on fin growth is acting at, or downstream of, mechanisms specifying size. Further, the phenotypic similarities between *alf* and those of FK506-treated fins raise the potential that these two mechanisms may be integrated.

**Figure 1.**
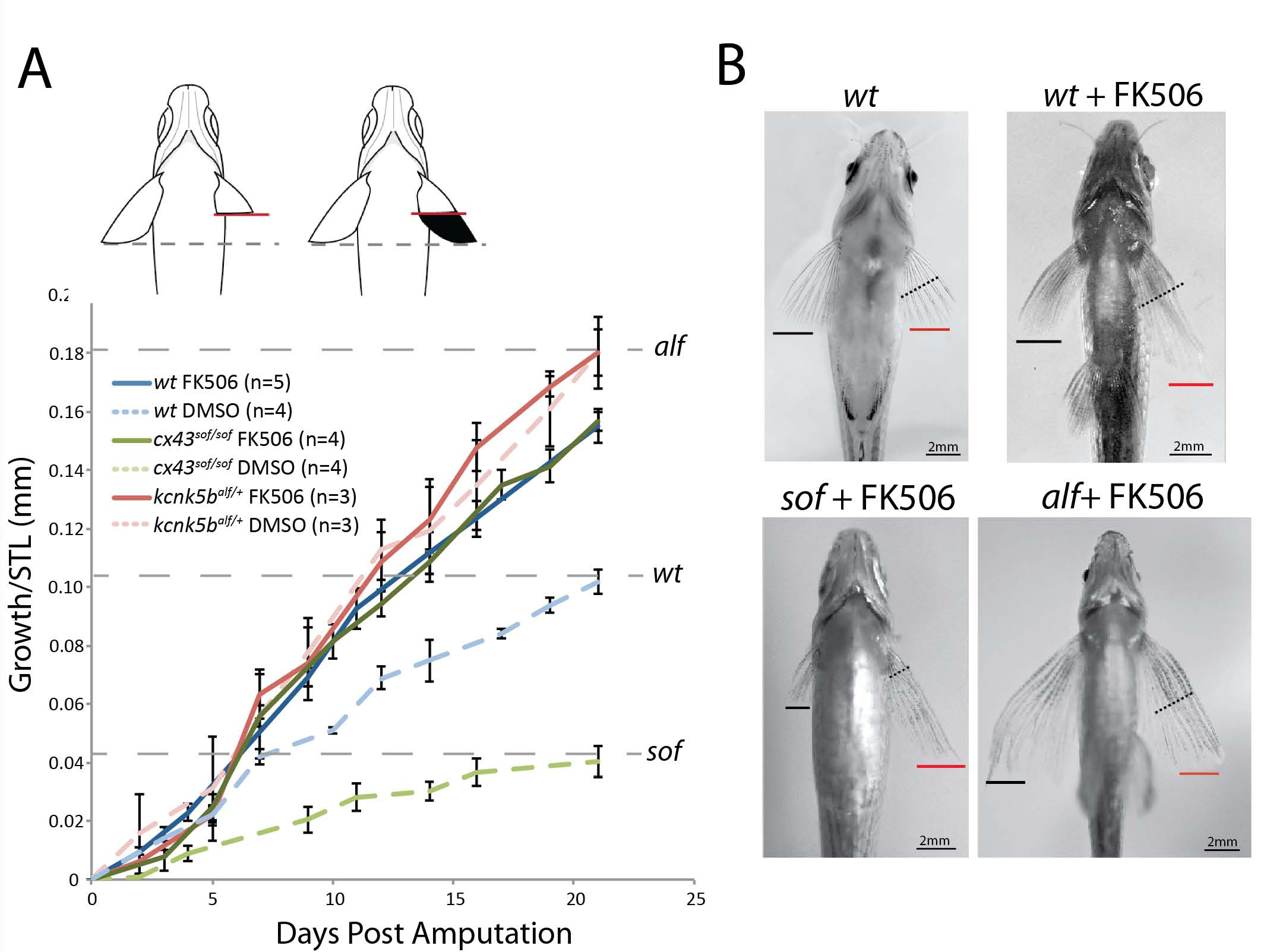
Calcineurin inhibition is sufficient to override genetic encoding of size. A) Regenerative growth of zebrafish pectoral fins after resection to ~50% of pre-cut fin length at 0 days post amputation in different genetic backgrounds with short- (*cx43^sof/sof^*) or long- (*kcnk5b^alf^*) fin size relative to wild-type fish. Dashed lines indicate pre-cut pectoral fin length. Data normalized to standard length (STL) of each fish. B) Representative ventral view of pectoral fins after regeneration. Black line indicates unoperated pectoral fin length. Red line indicates pectoral fin length after treatment with FK506 during regeneration. Black dashed line highlights site of amputation. Growth rate of FK506-treated regenerating caudal fins. D). Caudal fins were resected to 50% of their original size and treated with FK506. Growth rate is analyzed through 9 days post amputation, when regenerative growth rate for DMSO treated fins begins to slow. Two-tailed t-test p<0.0007 for suppression of growth in the absence of kcnk5b. Error bars represent ± SEM.

### Kcnk5b activity is critical for growth effects of FK506

Fish deficient for *kcnk5b* have normal fin proportions and growth (Perathoner et al., 2014). To assess the role of Kcnk5b in calcineurin growth regulation, we resected the pectoral fins of *kcnk5b^-^/-* fish and asked if the channel is needed for the FK506 growth response. Unexpectedly, the growth effects driven by FK506 treatment are suppressed in *kcnk5b^-^/-* fish (Figure 2A-C). This effect was seen in both pectoral fin as well as caudal fin regeneration (Figure 2D). These data reveal that Kcnk5b is a critical component of FK506-mediated regulation of growth and size. Some additional growth occurs after FK506 treatment in *kcnk5b*^-^/- fish, raising the possibility that residual function of the *kcnk5a* paralogue or other targets of FK506 may contribute in part to the FK506 response

**Figure 2.**
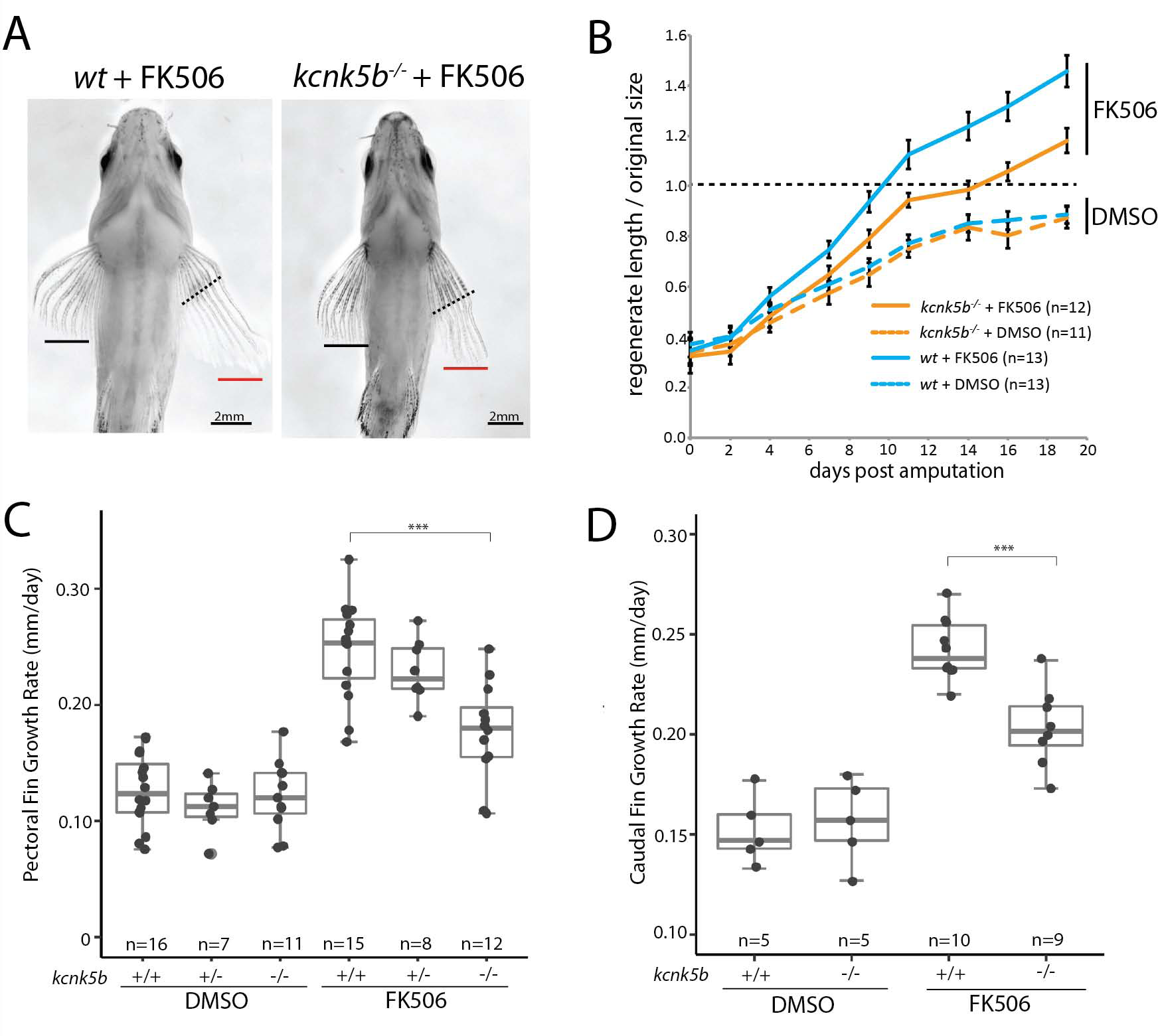
Kcnk5b mediates the effect of calcineurin in regulating proportion. A) Example of growth of pectoral fins of wildtype or *kcnk5b* deletion mutants regenerating after treatment with FK506. Dashed line indicates plane of section; black bar extent of growth of untreated fin; red, resected fin. B) Growth rates of wildtype and *kcnk5b* deficient zebrafish. Dashed line, size of original pre-cut fin; Error bars represent ± SEM. C) Growth rate of regenerating fins of different kcnk5b genotypes treated with FK506. Gray dots represent individual fish. Two-tailed t-test p<0.0002 for suppression of growth in the absence of *kcnk5b*. Data is from four independent experiments. (C) Representative ventral view of pectoral fins after regeneration.

### Modification of *kcnk5b* function by calcineurin

Kcnk5b could mediate FK506 growth effect indirectly or through modification of calcineurin regulation. Increasing the levels of wild-type *kcnk5b* locally in the fin is sufficient to induce fin overgrowth (Perathoner et al., 2014). As large changes in transcription are observed in regenerating fins treated with FK506 (Kujawski et al., 2014), we hypothesized that FK506 mediated overgrowth could be due to upregulation of *kcnk5b* expression levels. However, we find that expression of *kcnk5b* is not significantly altered in wild-type regenerating fins after FK506 treatment (Figure 3A). To test the potential for direct modulation of channel activity by FK506/calcineurin, we assessed the change in conductance of Kcnk5b channel variants in *Xenopus* oocytes. Oocytes expressing wild-type zebrafish Kcnk5b show decreased conductance when treated with FK506 (Figure 3D-E; see Nam et al., 2011). Similarly, treatment of oocytes expressing *kcnk5b* with VIVIT, an independent peptide inhibitor of calcineurin, showed a comparable decrease in conductance, supporting a role of calcineurin in regulation of channel activity (Figure 3E **inset**).

**Figure 3.**
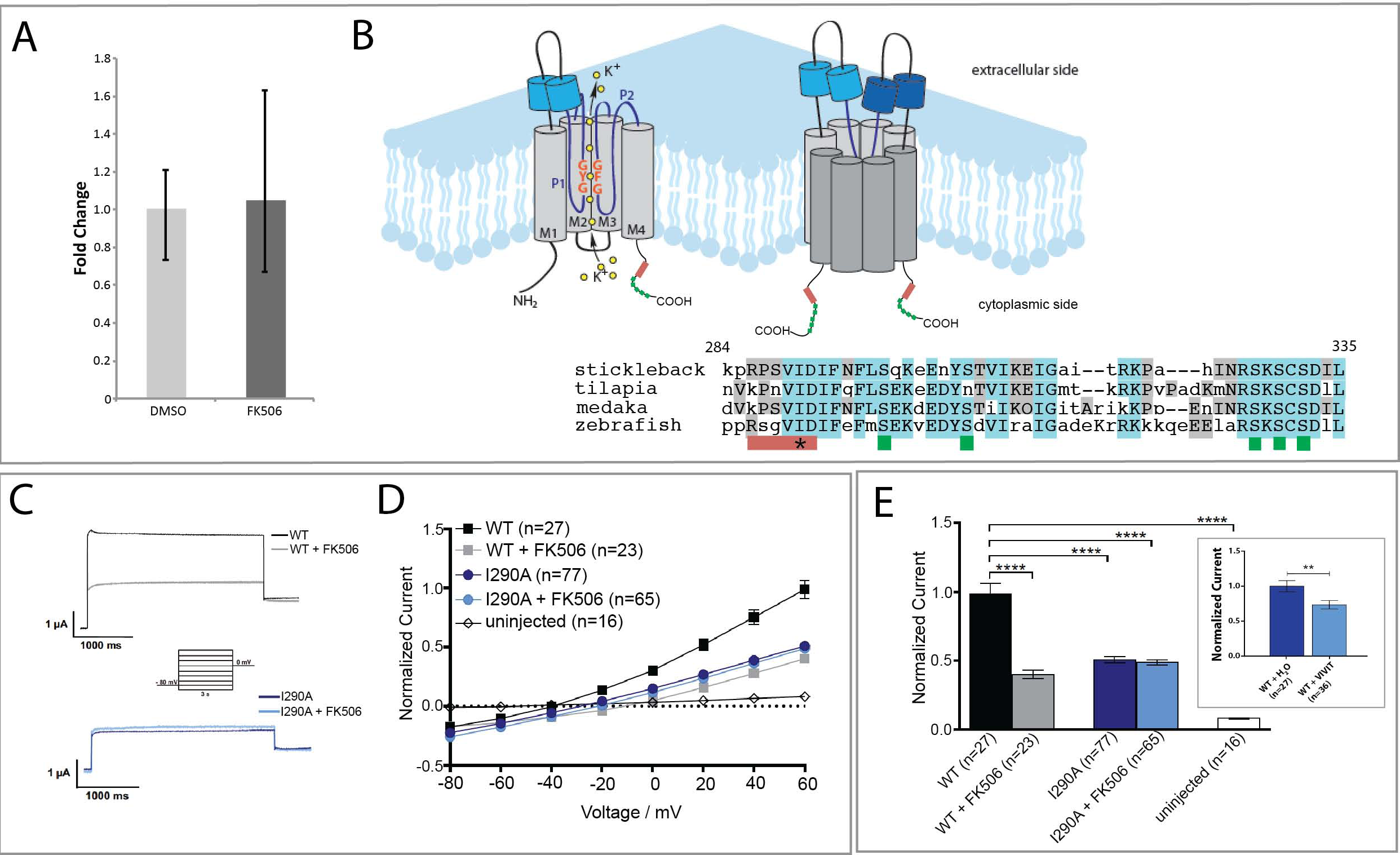
FK506 effect on Kcnk5b activity is modulated by the cytoplasmic domain of the channel. (A) qRT-PCR comparing relative levels of *kcnk5b* in FK506-treated, 5-day post-amputation pectoral fin blastemas to values of DMSO-treated controls (n=5, ± SEM). (B) Multiple sequence alignment of Kcnk5b protein highlighting putative calcineurin binding domain with resemblance to calcineurin binding in TRESK (red bar, PQIVID (Czirják and Enyedi, 2006) and a nearby suite of highly conserved serine residues (green bars). Asterisk indicates site of mutagenesis in (C-E) (I290A). Alignment generated by MUSCLE (Edgar, 2004). Diagram adapted from (Perathoner, 2013). (C) Representative electrophysiology current traces from voltage clamp recordings in *Xenopus* oocyte injected with wild-type or mutant I290A Kcnk5b cRNA and treated with FK506 or DMSO. The membrane potential was clamped at a reference potential of −80 mV and then varied by incremental 20mV steps to a range of −100mV to +60mV. (D) Current is suppressed in wild-type Kcnk5b homodimers through treatment with FK506 or by mutation of the calcineurin binding site (I290A). (E) Recorded currents were normalized to the value recorded for oocytes injected with the wild-type channel at +60 mV. *Inset*, Xenopus oocytes expressing wild-type zebrafish Kcnk5b were injected with VIVIT peptide or carrier. Patch clamp experiments registered the resulting conductance. ** p<0.001, n, number of oocytes assessed.

It has been shown that the activity of another KCNK two-pore potassium channel family member, Kcnk18/TRESK, is regulated by direct binding of calcineurin to the intracellular loop of the channel. This binding affects downstream dephosphorylation events on conserved serine residues on the C-terminus of the channel (Czirják and Enyedi, 2006; Czirják et al., 2004). Intriguingly, we identified a putative calcineurin binding site in the cytoplasmic C-terminus of zebrafish Kcnk5b that is similar to that identified in KCNK18/TRESK (Figure 3B). Czirják et al previously demonstrated that an isoleucine to alanine mutation in this domain in TRESK attenuated the effect of calcineurin and its binding to the channel (Czirják and Enyedi, 2006). We mutated the co-responding residue in the presumptive binding site in Kcnk5b (I290A; Figure 3B) and assessed the effect on conductance of the channel in oocytes. Oocytes expressing Kcnk5b/I290A had lower conductance than wildtype, but were similar to wild-type channels treated with FK506 (Figure 3C-E). Importantly, the Kcnk5b/I290A expressing oocytes were also non-responsive to FK506 treatment. Thus, the effect of the altering the molecular characteristics of this site on the Kcnk5b C-terminal intracellular domain is comparable in effect to chemical inhibition of calcineurin by FK506. This result suggests that, similar to KCNK18, the activity of Kcnk5b is modulated by calcineurin through interactions with the cytoplasmic C-terminus.

The *alf* mutation affects the last transmembrane domain of the channel located just prior to the predicted calcineurin binding site (Perathoner et al 2014, Figure S3). In an effort to screen for further variants affecting growth within this region, we used CRISPR-targeted Cas9 genetic editing with guides targeted to the fifth exon of *kcnk5b*, which contains the transmembrane and C-terminus, and screened adults for somatic clones sufficient to cause overgrowth. We recovered fish having overgrowth caused by guides targeting the transmembrane region of the channel. Analysis of Kcnk5b in these clones revealed formation of frameshift mutations leading to early termination (Figure S3). Thus, loss of the last transmembrane domain and the cytoplasmic tail of Kcnk5b is sufficient to lead to increased proportion.

### Calcineurin signaling does not re-specify positional cues, rather overrides them

We have demonstrated an interaction between Kcnk5b and calcineurin-mediated signaling in regulating growth. However, the ability of this module to specify growth or identity of the fin remains unclear. Kujawski *et al*. (2014) provide evidence of perdurance of proximal markers in treated fins and suggest that calcineurin inhibition leads to a re-specification of the regenerating tissue to a proximal identity. However, these data are also consistent with the hypothesis that the treatment leads to a sustained increase in the rate of growth that then is associated with broader domains of regional markers in the enlarged regenerated tissue. To address the effect of calcineurin on altering positional information of the fin, we mirrored classic experiments of Maden (1982) in regulation of positional identity of limb regenerates in the salamander and asked if limited alteration of calcineurin could respecify identity of the fin regenerate and thus its interpretation of fate.

We first approached this question by addressing how fins respond when FK506 is removed. Pectoral fins regenerating under the influence of FK506 exhibited increased growth (Figure 2, S1). However, after removal of the drug, either prior to achieving their original size, or in fins that had surpassed their original size, regenerative growth quickly stopped regardless of the extent of the fin regenerate (Figure 4A). Thus, FK506 induces an acceleration of growth that requires continual presence of the drug for sustained overgrowth. We extended these studies to address if the identity of the regenerated tissue was altered by treatment with FK506. As amputated fins will regenerate back to a size similar to that originally specified during development, the fin tissue remaining after amputation retains information concerning size and proportion that is then integrated in the regenerate. However, if overgrown fins caused by FK506 inhibition of calcineurin are resected, either into the original fin tissue or within a regenerated portion of the fin, these resulting regenerates grew only to the original size of the fin prior to treatment with FK506 (Figure 4B,C). Similarly, FK506-treated overgrown fins cut beyond the original wild-type fin length failed to grow further upon amputation without the presence of the drug (Figure 4C). Similar effects were seen in when tested in both pectoral as well as caudal fin regenerates (Figure S2). Thus, Kcnk5b/calcineurin signaling does not lead to lasting alterations in positional identity, but rather regulates rate of growth leading to enhanced fin proportions.

**Figure 4.**
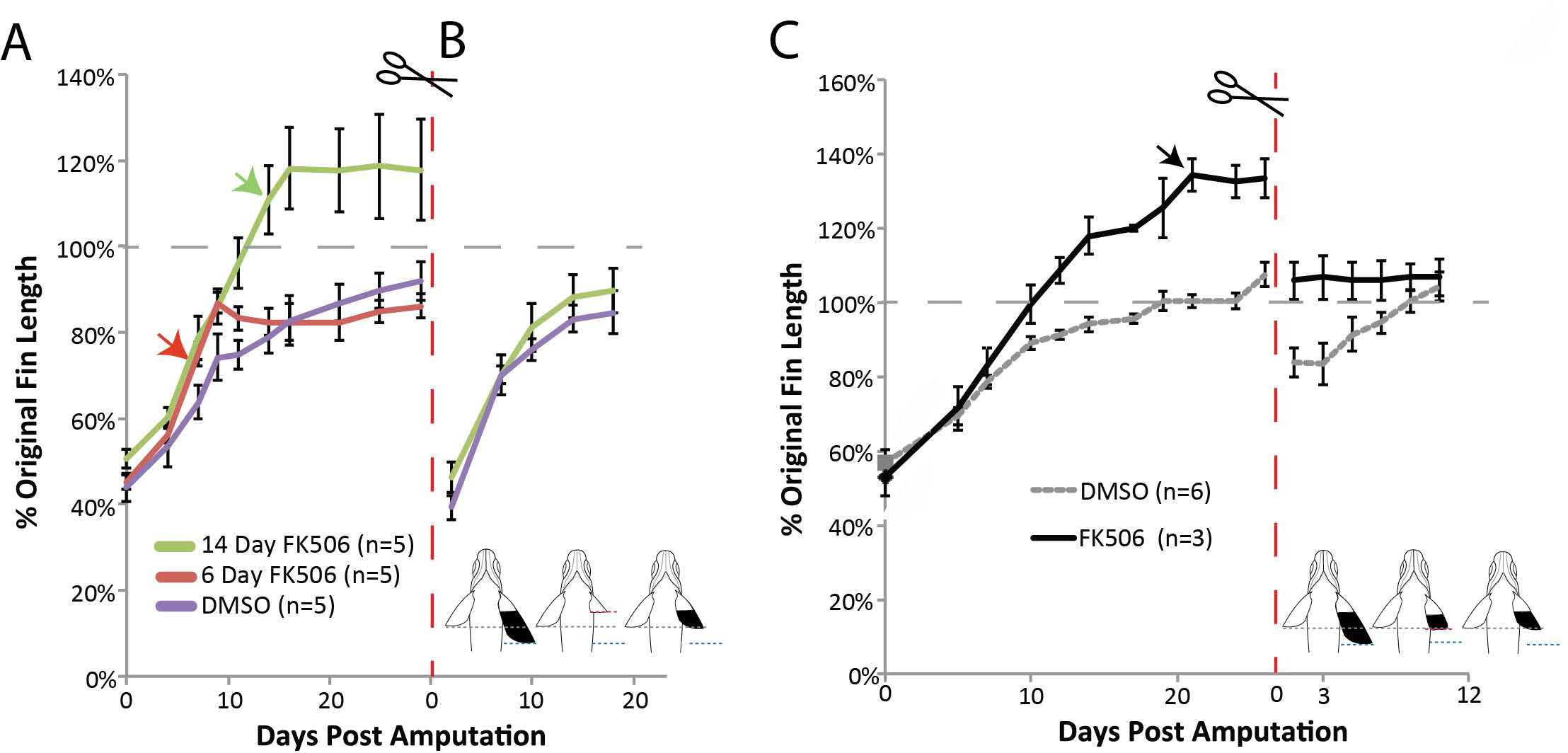
Calcineurin inhibition establishes regenerative growth rate independently of positional identity. A) Pulse of FK506 treatment for 6 or 14 days post-amputation to assess effects of transient inhibition of calcineurin by FK506 on growth and patterning of the fin. Fins were cut to 50% original length and allowed to regenerate in the presence of FK506. Arrows indicate end of drug treatment (n=5 fish per treatment group). (B) Subsequent regeneration of the pectoral fins from (A) in the absence of FK506 after resection at original cut site to determine if fin allometry was re-set to a larger size through previous FK506 treatment. (C) Second regeneration of FK506-treated fins cut at a site within the regenerated tissue demonstrating memory of original positional information and recovery of wild-type size within the pectoral fin. Gray dashed line indicates pre-cut fin length. Red dashed lines indicate second resection. Error bars represent ± SEM.

## Discussion

### Bioelectric integration of growth and form

Bioelectric potential is manifest as a resting voltage state of cells (*Vmem*). While it is clear that specialized cells such as nerves and cardiomyocytes have honed the regulation of electrical potential for specific functions, a broader role of this signaling in development and homeostasis is becoming apparent as it is necessary for normal pattern and growth of diverse organ systems (Bates, 2015; Beane et al., 2013; Dahal et al., 2012; Levin, 2014a; Levin, 2014b). Natural and experimental changes of resting potential can have broad effects in morphogenesis. A key property of bioelectric signaling is the capacity to coordinate responses across cells and tissues. Many tissues are electrically coupled through the action of gap junctions, and localized signals can expand via paracrine signaling by local fluctuations in electrical fields (Levin, 2012). Thus, the integration of cellular- and tissue-level responses to inductive signals as well as integration of patterning cues within developing organs can be orchestrated though electrical coupling of cells such that a unified structure is formed and maintained. Perturbations of such electrically coupled-developmental systems could be a factor for the coordinated transformations observed in evolution (Gould, 1966; Thompson, 1917) and broad patterning phenotype observed in some diseases (Bates, 2013; Dahal 2017). The role of *kcnk5b* in regulation of proportion was identified in genetic screens through its actions in causing altered scaling properties of adult fins (Perathoner et al 2014). We show that Kcnk5b is a key component to increased fin growth caused by calcineurin inhibition. Importantly, our work here demonstrates that Kcnk5b/calcineurin signaling is sufficient to increase growth regardless of innate size and genotype and thus override, but not re-specify, positional identity of the fin.

We demonstrate that Knck5b, like Kcnk18, can signal through calcineurin via a region in its C-terminus. Altering the presumptive binding site on the C-terminus of Kcnk5b (I290A) is sufficient to mirror the effect of calcineurin inhibition and to remove sensitivity to FK506 treatment for conductance. FK506 treatment of Kcnk5b in oocytes leads to decreased current. Similarly, the I290A mutation also mirrors this response. In contrast, *alf* mutants (Perathoner et al., 2014) show increased conductance in oocytes. This difference of the direction of conductance with similar growth outcomes may also point to the importance of signaling events downstream of changes in conductance that drive fin growth. The C-terminus of mouse Kcnk5/TASK-2 interacts directly with Gβγ subunit of heterotrimeric G-proteins (Añazco et al., 2013), and work on other two pore potassium channels, such as TREK-1 and TREK-2, has revealed dynamic regulation by cAMP/PKA, DAG/PIP/PKC and nitric oxide/cGMP/PKG pathways (Enyedi and Czirják, 2010). Modulation of Kcnk5b activity by conductance, or through interactions with its C-terminus, may act to adjust and/or respond to changes in voltage to regulate growth rate. Interestingly, we have found that premature truncation mutations leading to channels that lack the last transmembrane domain and the C-terminus of Kcnk5b result in fin overgrowth comparable to that seen in the *alf* mutant, which affects the identical region (F241Y**, Figure S3)**. As null alleles of *kcnk5b* do not cause an overgrowth phenotype, the truncation mutation appears to cause an increase in function of the channel. Alteration of the pore structure or conformation/presence of the C-terminus may cause dysregulated channel activity leading to overgrowth. The regulation of calcineurin may reflect downstream response to changes in conductance or conformation and thus, link calcium signaling to changes in pore conductance and/or conformation.

### Coordination of growth and proportion

Regulation of overgrowth by Kcnk5b/calcineurin signaling is coordinated among diverse tissues of the developing and regenerating fin to establish a larger, functional structure. This coordination may be attained through a re-interpretation of positional information within the fin such that regenerating tissues replace the missing components with tissue in register with the distal edge of the remaining fin stump. Positional information within the fin may be set up and driven by differential gene expression patterns in development (Rabinowitz et al., 2017; Tornini and Poss, 2014). Rabinowitz *et al*. (2017) detail expression profiles of wild-type fins that support dynamic regulation of gene expression across the proximal-distal aspect of the fin. Interestingly, a significant class of genes differentially isolated in these studies was ion channels, suggesting that bioelectric signaling may be a key factor of this asymmetry and establishment of positional cues. The work on calcineurin by Kujawski *et al*. (2014) also would point to calcineurin signaling as mediating position and being able to re-specify identity when altered. This mechanism was founded on extended gene expression domains in the proximal component of the fin and delayed distal bifurcation of the rays after calcineurin inhibition. However, an alternate explanation for the observed changes in pattern is that the expanded proximal molecular and anatomical identities observed in FK506-treated fins are a consequence of increased growth rate, such that larger regional domains, interpreted as position, are generated rather than specific changes in identity *per se*. This hypothesis would explain observed overgrowth as well as extended branching of the fin, but would be independent of changes in positional memory. Alternatively, positional information is independent of size instructive signals suggesting that modification of identity markers stemming from Kcnk5b/calcineurin signaling is coincident, but independent from, mechanisms of size determination and relative proportion.

Our data link the growth regulation of Kcnk5b and calcineurin and demonstrate that their function is not sufficient to re-specify identity, rather to override it. Establishing, or re-establishing, identity in the fin may require signals present only during development and may not be malleable during regeneration. Long-term treatment of fish during development with FK506 leads to global alterations in growth of the body that make comparison of proportion difficult (Figure S4). Thus, identification of these developmental cues to regulate proportion and size will require dissociation by other means. Our findings suggest that the regulation of growth by Kcnk5b/calcineurin fulfills the general requirements of a scaling property previously defined by relative growth comparisons and represented by the rate constant of Huxley and Teissier (1939). The encoding and specification of absolute size, however remains undefined.

## Acknowledgements

The authors wish to thank Drs. Christopher Antos and Satu Kujawski for their discussions of data on calcineurin prior to publication. *Sof* mutant lines were kindly provided by Dr. Kathryn Iovine. This work was supported in part by NSF GRFP and DDIG DEB-1407092 to JMD, and Grant HD084985 from NICHD, a John Simon Guggenheim Fellowship to MPH as well as support from the Children’s Orthopaedic Surgery Foundation at Boston Children’s Hospital.

---

**Supplemental Figure 1.**
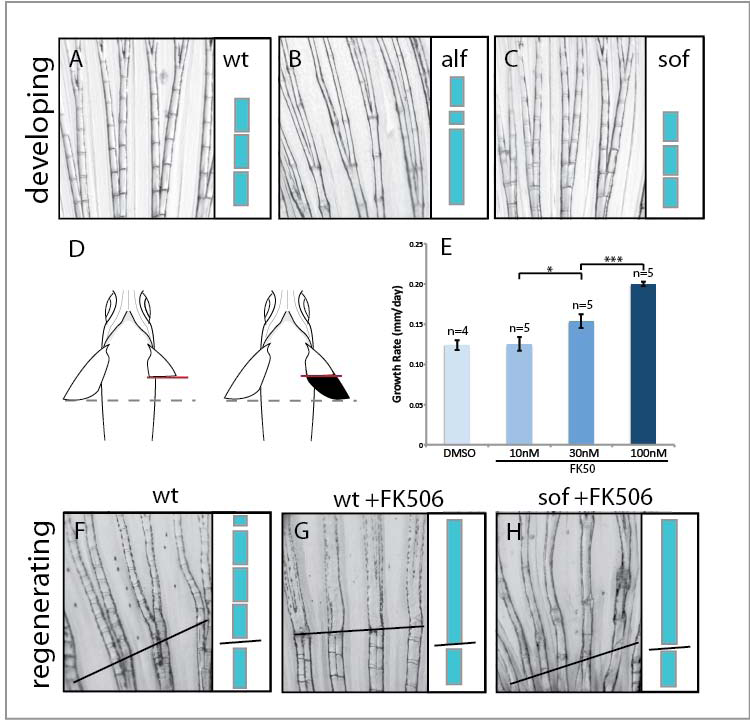
Proportion and patterning of zebrafish fins is enhanced by calcineurin inhibition. Fin growth occurs through sequential addition of a regular pattern of lepidotrichia hemi-ray dermal bone segments at growing end of the fin. Pattering of fin segmentation in A) wildtype, B) *alf* and C) *sof* pectoral fins; stacked blue rectangles model sequential addition and segmentation of individual rays. D) Schematic of regenerating pectoral fin assay to assess relative scaling. One pectoral fin is cut to approximately 50% of original size and allowed to regenerate. Comparisons to the contralateral side enables analysis of previous size and effect of treatment on non-regenerating fins. E) Dose response of FK506 treatment of pectoral fin growth. F-H) Pattern of segmentation of the lepidotrichia in F) regenerating wild-type, or regenerating FK506 treated wild-type or *sof* fins; solid line, plane of resection. While segments are normally restored during regeneration, regenerating fins treated with FK506 show elongated segmentation after regeneration similar to the *alf* phenotype (B). Error bars represent ± SEM. * p-value < 0.05. *** p-value < 0.001.

**Supplementary Figure 2.**
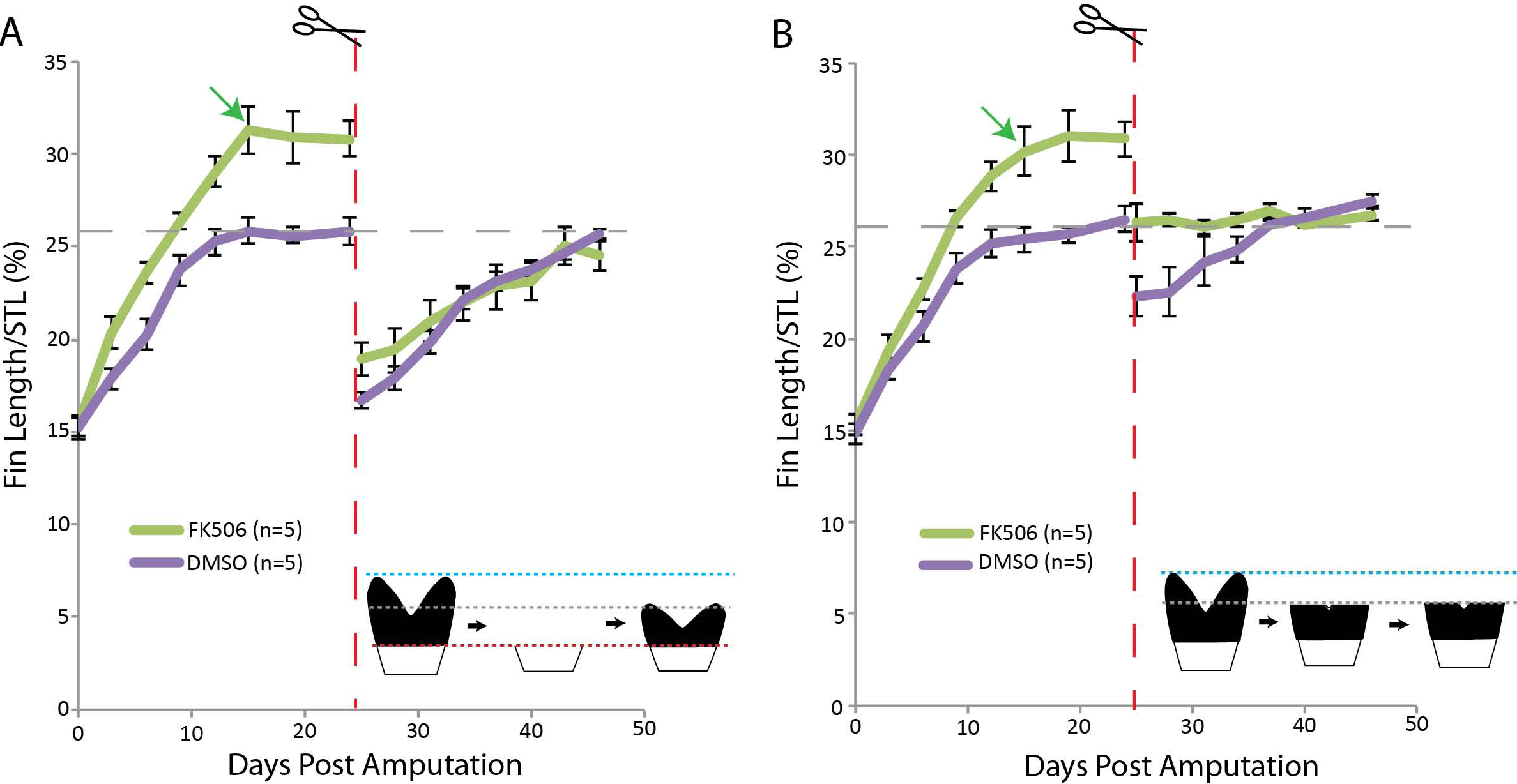
Calcineurin inhibition does not re-specify positional information of the caudal fin. Caudal fins were amputated and grown in FK506 for 15 days. The drug was removed (green arrow indicates stoppage of treatment) at which point the FK506 treated fins ceased further growth. These fins were then re-cut at either the original amputation plane (A) or at the site of the original fin length prior to FK506 treatment (B). Gray dashed line indicates precut fin length. Red dashed lines indicate second resection. Blue dashed lines indicate extent of FK506 overgrowth. Error bars represent ± SEM.

**Supplementary Figure 3.**
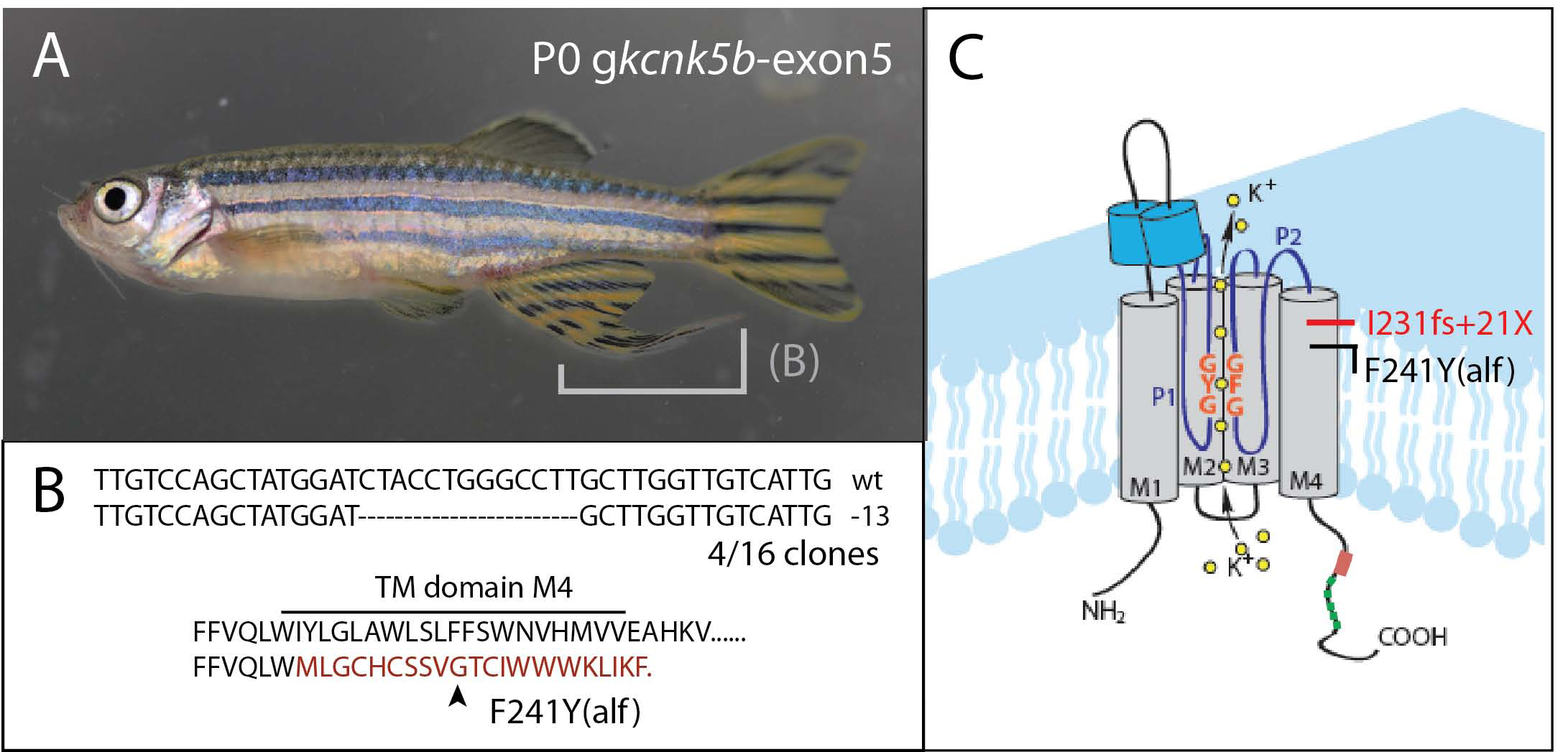
Deletion screen of *kcnk5b* reveals essential role of C-terminus. Using specific guide RNAs against the last exon of *knck5b* encoding the last transmembrane and cytoplasmic tail of the channel, we screened injected founders for evidence of overgrowth. We identified localized clones having specific overgrowth of the fins. B) Analysis of the changes in the overgrown tissues demonstrated presence of local deletions in *kcnk5b* predicted to cause truncation of the channel in a comparable location as to the *alf* mutation (C).

**Supplementary Figure 4.**
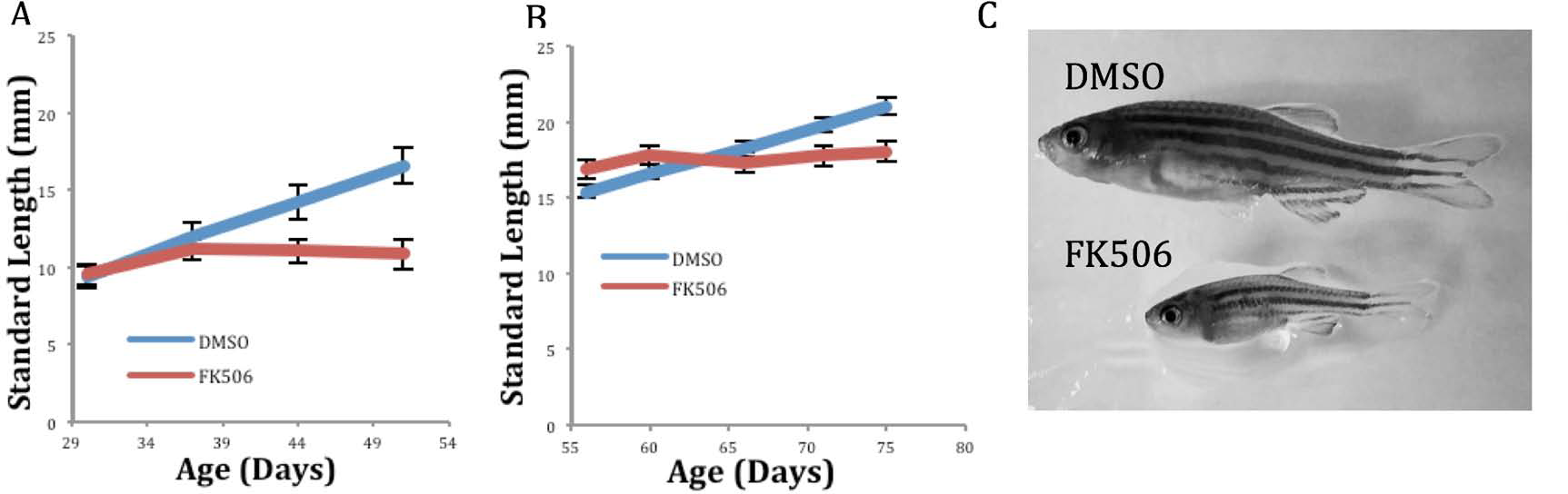
Growth deficit during FK506 treatment of juveniles. Wild-type juvenile zebrafish treated with FK506 did not grow when treated starting at ages of 30 days (A) or 55 days (B). C) Representative image of stunted growth in FK506 treated fish relative to DMSO treated siblings. n=10 fish per group. Error bars represent ± SEM.

## Supplemental Document 1. Experimental Procedures

### Fish Husbandry

The zebrafish AB strains and Tübingen (Tü) strains carrying *albino* mutation were used as background for all experiments. Fish were bred and maintained as previously described (Nüsslein-Volhard and Dahm, 2002). All experimental procedures involving fish conform to AAALAC standards and were approved by institutional IACUC committees. A complete description of the husbandry and environmental conditions in housing for the fish used in these experiments is available as a collection in protocols.io dx.doi.org/10.17504/protocols.io.mrjc54n For all experiments, adult stages defined by reproductively mature fish >3 months old were used for analysis. Mutant alleles used in this work are *alf ^dt30mh^*, *sof^dj7e2^*, and *kcnk5b^j131x8^*.

### Fin Regeneration and Measurements

Fish were anesthetized by treatment with tricaine for measurements and pectoral fin amputation. Unless otherwise indicated, pectoral fins of wildtype and mutants were resected to roughly 50% their pre-cut length using standard surgical scissors. Fin length was measured before, during and after regeneration using handheld calipers and measurements were normalized to the fish standard length, determined as the length from the tip of the snout to the posterior end of the caudal peduncle.

### FK506 Treatment

Stock FK506 (Sigma) was dissolved in dimethyl sulfoxide (DMSO; Sigma). Fish water was treated with 100nM FK506 unless otherwise indicated. Up to five individual zebrafish were housed in 1L of FK506- or DMSO-treated fish water and were fed daily with live artemia. Water and drug were refreshed every other day throughout the duration of the experiment.

### qRT-PCR

Five days post amputation, pectoral fin regenerates were collected from five individuals of each treatment group. For RNA extraction, tissue was homogenized in TRIzol Reagent (Invitrogen). cDNA synthesis was performed with oligo dT primers using SuperScript III Reverse Transcription Kit (Invitrogen). qRT-PCR was performed with Power SYBR Green Master Mix (Applied Biosystems) on Applied Biosystems ViiA 7 Real Time PCR System. Cycling conditions: 10 minutes at 95°C; 40 cycles of 15 seconds at 95°C followed by 1 minute at 60°C; melting curve analysis with 15 seconds at 95°C, 1 minute at 60°C and 15 seconds at 95°C. Temperature was varied at 1.6°C/s. Expression levels were normalized relative to β-actin. ddCt was used to calculate fold change in FK506-treated relative to the average of DMSO-treated fish. *Kcnk5b* primers: 5’-TTGTAGCCGTCTGTGACCAA-3’, 5’-AGTACCGCACCCAAACTGTC-3’. *β-actin* primers: 5’-CAACAACCTGCTGGGCAAA-3’, 5’-GCGTCGATGTCGAAGGTCA-3’.

### Electrophysiology

The cDNA of Kcnk5b was subcloned into pSGEM for cRNA expression. The plasmid was linearized using *NheI* and cRNA was *in vitro* synthesized using the T7 mMessage mMachine kit (Ambion). *Xenopus laevis* oocytes were provided by Ecocyte Bioscience. Oocyte handling, injection and electrophysiological recordings were as previously described(Seebohm et al., 2005). Briefly, stage V oocytes were injected with 4 ng of cRNA encoding wildtype or mutant I290A *kcnk5b* and stored for 3 days at 18°C in Barth’s solution containing (in mmol/L): 88 NaCl, 1.0 KCl, 2.4 NaHCO_3_, 0.33 Ca(NO_3_)_2_, 0.4 CaCl_2_, 0.8 MgSO_4_, 5 Tris-HCl, penicillin-G (63 mg/L), gentamicin (100 mg/L), streptomycin sulfate (40 mg/L), theophylline (80 mg/L); pH 7.6. Two-electrode-voltage--clamp recordings were performed at 22°C using a Turbo Tec-10CD amplifier (NPI electronics), Digidata 1322A AD/DA-interface and pCLAMP 9.0 software (Axon Instruments Inc. / Molecular Devices). Before measurement oocytes were pre-incubated in 0.5 % DMSO (control) or in 50 µM FK506 (InvivoGen) for 1 h at 18°C in ND96 recording solution containing (in mM): 96 NaCl, 4 KCl, 1.8 MgCl_2_, 1.0 CaCl_2_, 5 HEPES; pH 7.6. FK506 was dissolved in DMSO and FK506 dilution in ND96 was prepared freshly before every experiment. Recording pipettes were filled with 3 M KCl and had resistances of 0.5-1.5 MΩ. Data were analyzed with Clampfit 9.0 (Molecular Devices Corporation), Excel (Microsoft) and Prism6 (GraphPad Software). The Kcnk5b I290A variant was generated by QuikChange II (Agilent). Mutagenesis primers : 5’-CTCGCTCTGGAGTCGCTGACATCTTTGAG-3’, 5’-CTCAAAGATGTCAGCGACTCCAGAGCGAG-3’.

### CRISPR generation of *kcnk5b* somatic clones

The fifth and terminal exon of zebrafish *kcnk5b* encodes the last transmembrane domain and cytoplasmic C-terminus. We designed guide RNAs (gRNA) tiling this exon to screen for sufficiency of deletions in this region to affect fin growth in mosaic injected animals. The ChopChop online was used to design gRNAs to limit predicted off-target gRNA cutting (Labun et al., 2016; Montague et al., 2014). The pool of 5 distinct gRNAs were simultaneously injected to blanket the start of the exon. These gRNAs were assembled according to Gangon *et al*. (Gagnon et al., 2014). Briefly, oligos containing the gRNA target were annealing to a universal oligo containing the tracrRNA and SP6 promoter. The annealed oligo ends were then filled in with T4 polymerase for 20 minutes at 12˚C. gRNA was synthesized from this oligo template using Ambion MEGAscript SP6 Kit. For transcription efficiency, the first two bases of each gRNA were changed to ‘GG’ as there is evidence that these bases have less effect on Cas9 cutting efficiency or off-target binding than mutations closer to the PAM site (Fu et al., 2013; Hwang et al., 2013a; Hwang et al., 2013b). gRNAs were injected into single cell zebrafish embryos at a concentration of 50ng/µl blanket and 300ng/µl Cas9. To screen for deletion efficiency, the target exon was amplified from pools of three 24 hour embryos and the resulting amplicons were heated to 95°C and cooled at −0.1°C per second to form heteroduplexes. Following heteroduplex PCR, a T7 endonuclease digestion for 30 minutes at 37°C in NEB Buffer 2 was used to generate deletions in the presence of Cas9-induced indels. Primers for colony PCR cloning of the exon 5 screen: 5’-GGCAAATCAAACTGGTTAGTCC-3’, 5’-CGCTGTAGTCCTCGACCTTC-3’. gRNA sequences: 5’-ATCACTTTGTTTCCATACAG-3’, 5’-GACAAAGAATCTGTAGAGAG-3’, 5’-GGATCTACCTGGGCCTTGCT-3’, 5’-TTGGAACGTGCATATGGTGG-3’, 5’-CGTCATCGGTGGGCAGCCTG-3’.

